# RNA-associated glycoconjugates highlight potential ambiguities in glycoRNA analysis

**DOI:** 10.1101/2024.03.12.584655

**Authors:** Sungchul Kim, Yong-geun Choi, Kirsten Janssen, Jan-Willem H. Langenbach, Daan J. van den Brink, Christian Büll, Bhagyashree S. Joshi, Adam Pomorski, Vered Raz, Marvin E. Tanenbaum, Pascal Miesen, Zeshi Li, Chirlmin Joo

## Abstract

A recent ground-breaking study suggested that small RNA from mammalian cells can undergo N-glycan modifications (termed glycoRNA) ^1^. The discovery relied upon a metabolic glycan labeling strategy in combination with commonly used phase-separation-based RNA isolation. Following the reported procedure, we likewise identified an N-glycosylated species in the RNA fraction. However, our results suggest that the reported RNase sensitivity of the glycosylated species depends on the specific RNA purification method. This suggests the possibility of co-purifying unexpected RNase-insensitive N-glycoconjugates during glycoRNA isolation. The co-existence of two independent, yet highly similar molecular entities, complicates biochemical assays on glycoRNA, and calls for more specific approaches for glycoRNA analysis. To address this, we propose a control experiment that can help distinguish genuine glycoRNA species from co-purified glycoconjugates.

## Introduction

N-linked glycosylation is a major post-translational modification that affects folding, stability, and other cellular functions of secretory and membrane-associated proteins ^2^. N-glycosylation starts in endoplasmic reticulum by the assembly of high-mannose glycans and the transfer thereof to nascent peptides. The glycan is then trimmed by ER mannosidases and is further elaborated in the Golgi apparatus, where multi-antennary branching and extension, fucosylation, and sialylation are introduced^3^. Expanding the world of N-glycosylation, a recent study reported that specific small non-coding RNA species in mammalian cells are modified with sialylated and fucosylated N-glycans ^1^. These molecules, referred to as glycoRNA, were reported to localize to the surface of mammalian cells and were shown to interact with either specific Siglec family receptors or P-selectin ^1,4,5^.

Bioorthogonal labeling of glycans present in RNA isolates was a critical step in the discovery of glycoRNA ^1^. Carbohydrate analogs or their precursors modified with bioorthogonal chemistry tags, called metabolic chemical reporters (MCRs), are important tools in glycoscience, and enable the delineation of the biogenesis and function of glycosylation. MCR, combined with bioorthogonal labelling, is a widely-used method due to its simplicity of implementation, the rapidity of the chemical reactions, as well as the bio-compatibility and high specificity in biological environments ^6^. MCRs exploit the tolerance of cellular glycan biosynthetic pathways towards unnatural modifications, which are eventually incorporated into glycans^7^. For example, peracetylated N-azidoacetyl-mannosamine (Ac_4_ManNAz) has been used as MCRs for sialic acid labeling. Ac_4_ManNAz is converted into azido-sialic acids (SiaNAz) and is normally incorporated at the terminus of glycans ^8^. The azide tag in sialoglycans can be conjugated to alkyne-containing molecules, for instance, fluorophores or biotin, via copper-catalyzed azide– alkyne cycloaddition (CuAAC) or copper-free strain-promoted alkyne-azide cycloaddition (SPAAC) ^9,10^. The latter reaction was used to demonstrate the presence of glycoRNA ^1^.

A common method to obtain cellular RNA is to use acidic guanidinium-thiocyanate-phenol-chloroform (AGPC) phase partition ^1^. This technique relies on chaotropic agents for cell lysis and protein denaturation, followed by phenol-chloroform-based phase separation for isolation of RNA from other cellular components ^11,12^. Following alcohol precipitation of the RNA from the AGPC aqueous phase, the sample normally undergoes further clean-ups using proteases and DNases, after which the RNA are assumed to have high purity^12^. Silica solid-phase extraction has been a facile complementary method to AGPC-based RNA purification, as it is believed to yield RNA with exquisite purity ^13^. Although functionally pure RNA sample can be obtained, both methods rely upon the physical chemical properties of RNA. As a result, biomolecules with similar properties can be co-isolated^14,15^.

Here, we report that non-RNA N-glycoconjugates persist throughout RNA sample processing steps. We have found that the N-glycoconjugates are associated with RNA independent of extraction methods, including acidic phenol-chloroform and silica-based column extraction. In most biochemical assays, these N-glycoconjugates exhibit properties difficult to distinguish from what has been described for glycoRNA,^1,4,5^ in terms of enzymatic sensitivities and mobility in gel electrophoresis. A key difference between the N-glycoconjugates from glycoRNA is that the former is resistant to RNase digestion. However, we found that by a silica column clean-up after RNase treatment, as was performed in all reported procedure for glycoRNA preparation,^1,4,5^ the signals for the N-glycoconjugates were indeed lost in fluorescent gels or blots, apparently resembling the effect of RNase digestion of glycoRNA. The signals of the N-glycoconjugates can be rescued either by increasing alcohol concentration in buffers for silica column loading, or by an addition of exogenous RNA prior to loading into the column. We also demonstrate that the covalent attachment of glycans does not confer RNA the resistance towards RNase digestion, supporting the non-RNA nature of the N-glycoconjugates. Taken together, our data demonstrates RNA isolation methods are susceptible to the contamination of specific glycoconjugates. The potential co-existence of glycoRNAs with unknown N-glycoconjugates can cause complications in biochemical assays focused on glycoRNAs. Our conclusion is based on experiments performed in four laboratories using reagents that were independently purchased.

## Results

### Metabolically labeled glycoconjugates co-purified with RNA are RNase-resistant

The reported workflow for glycoRNA isolation starts by supplementing the cell culture with Ac_4_ManNAz, followed by extracting the total RNA using TRIzol, cleaning up RNA samples using multiple enzymes, and finally conjugating the SiaNAz-containing entities such as glycoRNA with dibenzocyclooctyne-biotin (DBCO-biotin). The samples were visualized on a biotin blot to check for the presence of bands with high apparent molecular weight as a readout for successful glycoRNA isolation^1^. In an attempt to develop an independent and efficient workflow for glycoRNA preparation, we employed a practical approach that employs mainly AGPC phase separation followed by ethanol precipitation in between processing steps and relies on labelling SiaNAz-containing molecules with dibenzocyclooctyne-Cyanine5 (DBCO-Cy5) (**Fig. 1a** and **Extended Data Fig. 1a**). This modified workflow enabled direct in-gel detection of RNA extracted from Ac_4_ManNAz-treated cells after separation by gel electrophoresis, eliminating the need for RNA membrane transfer. As expected, no fluorescent signal was observed in RNA obtained from DMSO-treated control cells (**Fig. 1b**). Importantly, the in-gel fluorescence band pattern was similar to the one observed after transfer to nylon or nitrocellulose membranes, validating that the visualization of the fluorescently labeled glycoconjugates is not affected by the blotting membrane (**Extended Data Fig. 1b**). We noted that signals from the labeled glycans were only stronger after transferring to nylon membranes compared to nitrocellulose membranes. Nonetheless, due to its practicality, we used direct in-gel Cy5 detection in further experiments.

**Figure 1.**
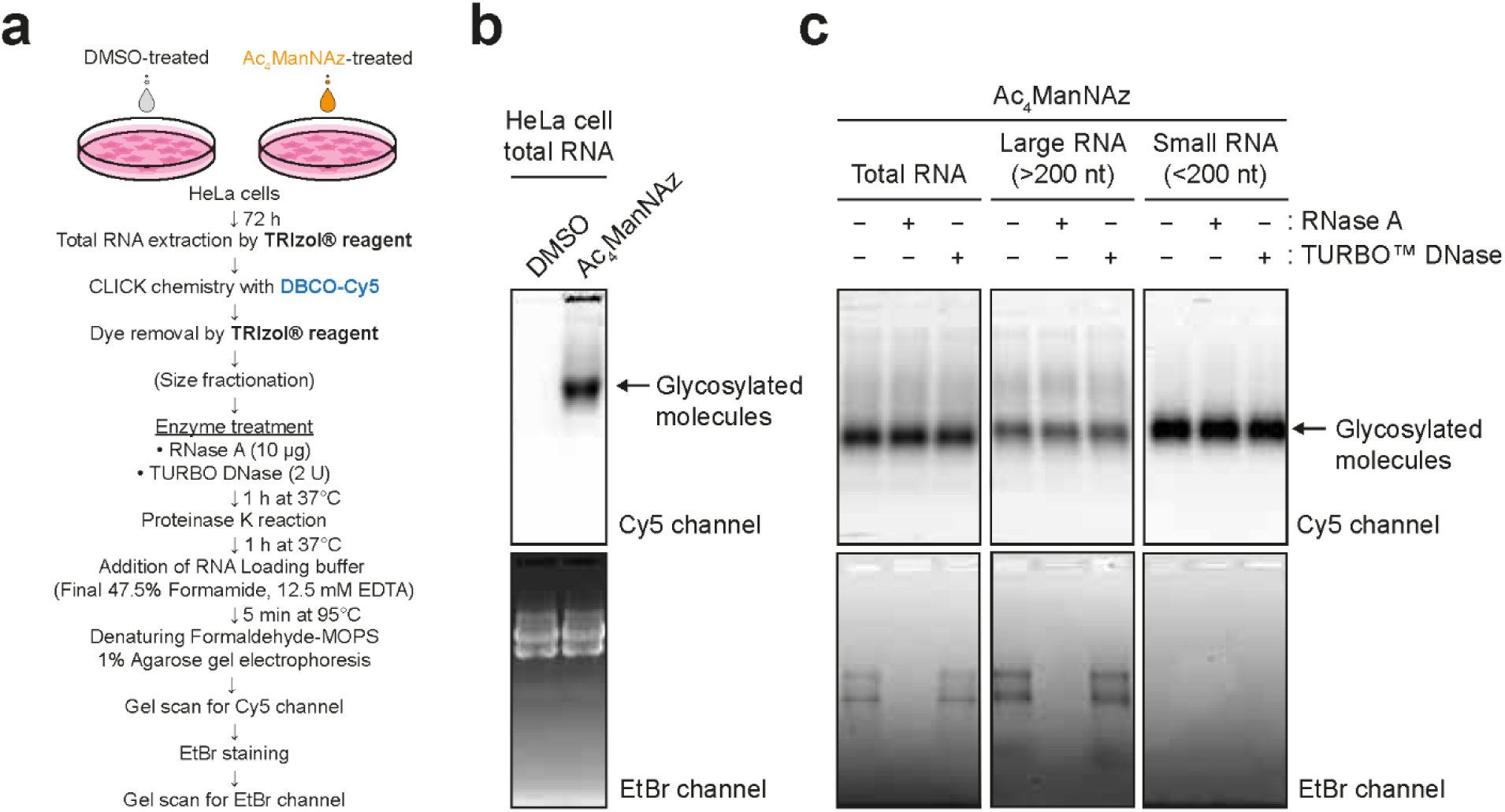
Glycoconjugates co-purified with RNA are insensitive to RNase treatment. **a.** Schematics of the experimental procedure for labeling and visualization. **b.** Glycan detection in Cy5 channel from a denaturing agarose gel without any enzymatic treatment condition. Dimethyl sulfoxide (DMSO) was used as an Ac_4_ManNAz untreated control. Ethidium bromide (EtBr) channel imaged on the gel scanner shows RNAs were intact. **c.** Glycan visualization in total RNA from HeLa cells or size-fractionated RNAs.

We then employed two checkpoint experiments to examine if we had isolated glycoRNA using the procedure above: RNA size fractionation and RNase digestion^1^. The previous reports demonstrate glycoRNA should be present in small RNA fraction (< 200 nt) when using silica column to fractionate the extracted total RNA. In line with the previous reports, the glycan signal was more intense in small RNA than in large RNA fractions, suggesting that the glycosylated moiety mainly coincide with small RNA (**Fig. 1c**). However, we observed that the glycan signal was not affected by either RNase or DNase treatments (**Fig. 1c** and **Extended Data Fig. 1c**), despite the similar electrophoretic properties to glycoRNA. At this point, via our procedure, a nuclease-resistant glycoconjugate(s) has been isolated.

### Specific sample processing steps lead to apparent RNase sensitivity of the glycoconjugates

We were intrigued by the electrophoretic similarities between the RNA-associated glycoconjugates and glycoRNA, and asked to what degree the RNA sample processing would be prone to contamination by the glycoconjugates. This is a critical consideration in the field because such similarity can potentially confound assays on glycoRNA^1,5^. We noticed that the RNA sample processing for glycoRNA detection requires multiple rounds of silica column-based purification and has a strict order for processing steps. We, therefore, set out investigating whether modifying the procedure could influence the recovery of the RNase-resistant glycoconjugates. In our modified sample processing procedure, we minimized the use of silica columns and instead used TRIzol phase separation followed by ethanol precipitation throughout initial RNA purification, dye removal, and clean-up after enzymatic reactions. Furthermore, we have modified the order of experimental steps, performing the click chemistry before enzymatic treatments (“early-click” procedure; **Fig. 2a**). In comparison, in the reported procedure, all RNA purification steps involved silica columns, and the click chemistry reaction was performed after enzymatic treatments. Finally, the RNA was again column-purified before gel electrophoresis and visualization (“late-click” procedure; **Fig. 2a**).

**Figure 2.**
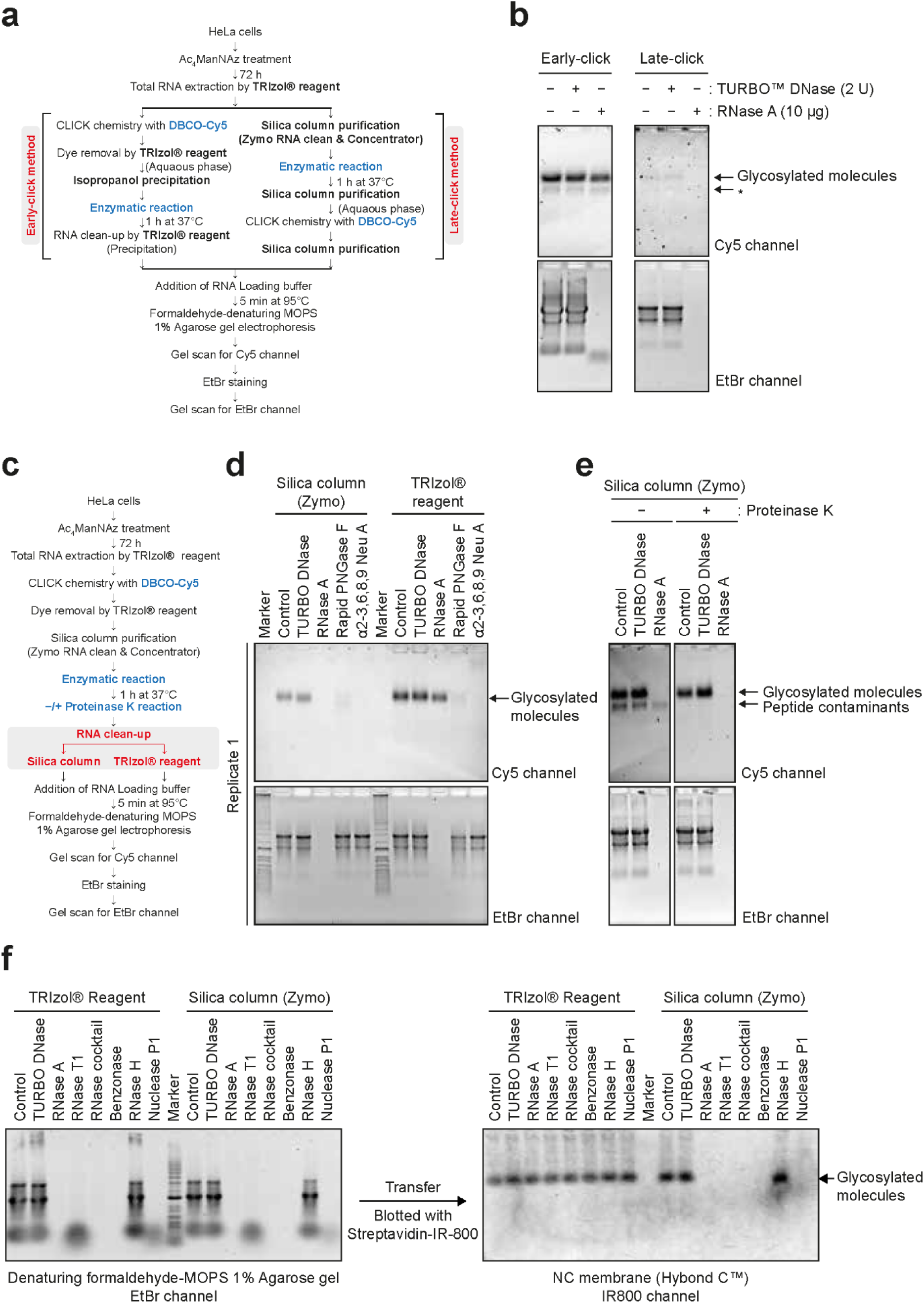
RNase sensitivity of glycosylated molecules depends on the RNA extraction method. **a.** Schematic comparison of early-click and late-click protocols. **b.** Glycan detection by early-click and late-click methods. The asterisk indicates the presence of labeled peptide contaminants (see panel **e**) **c.** Schematic representation of experiments in **Fig. 2d** and **2e**. **d.** Comparison of RNA clean-up steps between silica column and TRIzol protocols for glycan visualization. DBCO-Cy5 labeled RNA samples were reacted with TURBO DNase or RNase A or Rapid PNGase F or α2-3,6,8,9 neuraminidase A. Data represent one of three replicates. **e.** Glycan visualization with or without proteinase K treatment. **f.** Blotting of biotinylated glycosylated molecules treated with various nucleases using streptavidin-conjugated IR800 dyes on the nitrocellulose membrane. Right, gel image of EtBr-stained RNA samples. Left, fluorescent image using IR800-streptavidin in 800-nm channel.

We have compared the early-click and late-click procedures regarding their efficiency to recover the glycoconjugate. With the early-click approach, we observed strong fluorescent bands of high molecular weight in RNA gel electrophoresis, and a corresponding higher RNA isolative yield as indicated by the EtBr signal. These bands were insensitive to RNase (**Fig. 2b**, top left), while the total RNA was efficiently digested by RNase (**Fig. 2b**, bottom left). The late-click procedure yielded bands of the same apparent molecular weight but with a much weaker fluorescent signal, and correspondingly less total RNA. Notably, in the late-click procedure as was employed in previous reports, when the sample was treated with RNase, a loss of the glycan signal in agarose gel was observed. Our results indicate although the glycoconjugate persists throughout RNA sample processing, even after RNA degradation, the choice of the RNA purification method can critically influence the outcome. The silica column purification showed an overall lower recovery of the glycoconjugate and resulted in an apparent RNase sensitivity (**Fig. 2b**, top and bottom right, respectively).

To test the hypothesis that the final silica column had been the cause for the apparent RNase sensitivity, we took the early-click approach for its high RNA yields and kept all processing steps identical except for the last RNA purification (**Fig. 2c**), such that it would be a fair comparison between silica column and TRIzol followed by ethanol precipitation. Consistent with the results above, we found that the recovery of the glycan-labeled molecules was substantially poorer when silica columns were used to clean up the sample. While the RNase activity was confirmed by the degradation of total RNA (EtBr channel), the loss of fluorescent glycan signal upon RNase-treatment was only observed when silica column-based purification was used (**Fig. 2d** and **Extended Data Fig. 2a** and **b**). In previous reports, the loss of glycan signal has been used as a key indicator for a successful isolation of glycoRNA. Here we have shown the routinely used silica column in glycoRNA purification can cause an apparent RNase sensitivity for an RNase-resistant glycoconjugate.

### The RNA-associated glycoconjugate is modified by N-glycan but is not an RNA molecule

We further investigated the reactivity of the RNase-resistant glycoconjugate towards a wider array of enzymes. We observe that, regardless of whether RNA was purified with silica columns or TRIzol extraction followed by ethanol precipitation, the glycan signal was sensitive to the treatment with PNGase F and α2-3,6,8,9 neuraminidase A (**Fig. 2d**, and **Extended Data Fig. 2a** and **b**). The former is an amidase that cleaves oligomannose-, hybrid-, and complex-type N-glycans from glycoproteins/-peptides ^16^ and the latter is an exoglycosidase that removes terminal sialic acid residues linked to a penultimate sugar ^17^. The glycan signal was insensitive to high-amount proteinase K (20 μg per reation) treatment, although adding proteinase K resulted in the loss of additional bands appearing below the glycoconjugate bands (presumably peptide contaminants) in the gel image (**Fig. 2e**). Our results suggest that the detected glycosylated molecules in our experiments contain hybrid or complex N-glycans and exhibit the highly similar reactivities towards the digesting enzymes as reported for glycoRNA ^1^.

The assays described above have largely relied on RNase A and/or T1. To exclude that RNase resistance of the N-glycoconjugate had been due to the enzyme’s substrate specificity, we performed digestion assays on the glycan labeled molecule with an expanded collection of nucleases. While RNase A catalyzes the cleavage of single-stranded RNA (ssRNA) after pyrimidine nucleotides ^18^, RNase T1 specifically degrades ssRNA at G residues ^19^; benzonase can degrade various forms of DNA and RNA ^20^; RNase H cleaves RNA in RNA:DNA hybrids ^21^; and nuclease P1 hydrolyzes phosphodiester bonds in RNA and ssDNA without base specificity ^22^. Our results demonstrated that RNase cocktail, comprising RNase A and T1, and benzonase degraded RNA completely, and RNase T1 alone and nuclease P1 digested almost all RNA into small pieces, whereas RNase H did not result in RNA degradation (**Fig. 2f** and **Extended Data Fig. 2c**). Under all these conditions, signals corresponding to the N-glycoconjugate remained after TRIzol and the ensuing ethanol precipitation clean-up. The resistance of the N-glycoconjugate towards multiple nucleases supports it is not a nucleic acid.

In contrast, with silica column as the final clean-up step, the N-glycoconjugate signal disappeared in all samples with degraded total RNA (**Extended Data Fig. 2c**). To rule out any potential artifacts arising from the use of the Cy5 dye in in-gel fluorescence assays, we performed the click reaction with DBCO-PEG_4_-biotin to label metabolically azide-functionalized-glycans, as was employed in the previous studies^1,5^ (**Fig. 2f** and **Extended Data Fig. 2d**). We made the same observation as in the in-gel fluorescence experiments. Taken together, the association with the intact RNA appears to be crucial for the recovery of the N-glycoconjugate over a silica column, but the presence RNA is not required when the same molecule is centrifugally precipitated.

Currently, the only source to obtain glycoRNA has been limited to the isolation from cells, in combination with glycan labeling. It has not been possible to build the molecule in vitro, although a recent publication has provided structural indications for the chemical linkage between N-glycans and RNA^23^. To demonstrate the non-RNA nature of the N-glycoconjugates, we sought to make an artificially N-glycosylated RNA (neo-glycoRNA) to compare its enzymatic reactivities and electrophoretic properties with the N-glycoconjugates. To assemble the neo-glycoRNA, we took a photochemical approach^24^ to incorporate a strained alkyne (DBCO) to the guanosine residues of extracted cellular total RNA fragmented to the size of small RNAs (∼200 nt). We employed an azide-containing N-glycan^25^ (**Extended Data Fig. 3a**) fluorescently labeled at the sialic acid residue by a sialyltransferase.^26^ The N-glycan-azide was then clicked with the DBCO-containing RNA to afford the neo-glycoRNA (**Fig.3a**). We expect such entity to be structurally analogous to glycoRNA (**Extended Data Fig. 3b**), especially as guanosine had been proposed as an N-glycan attachment site^27^, although the modified uridine (acp^3^U) was experimentally supported to carry N-glycans^23^.

**Figure 3.**
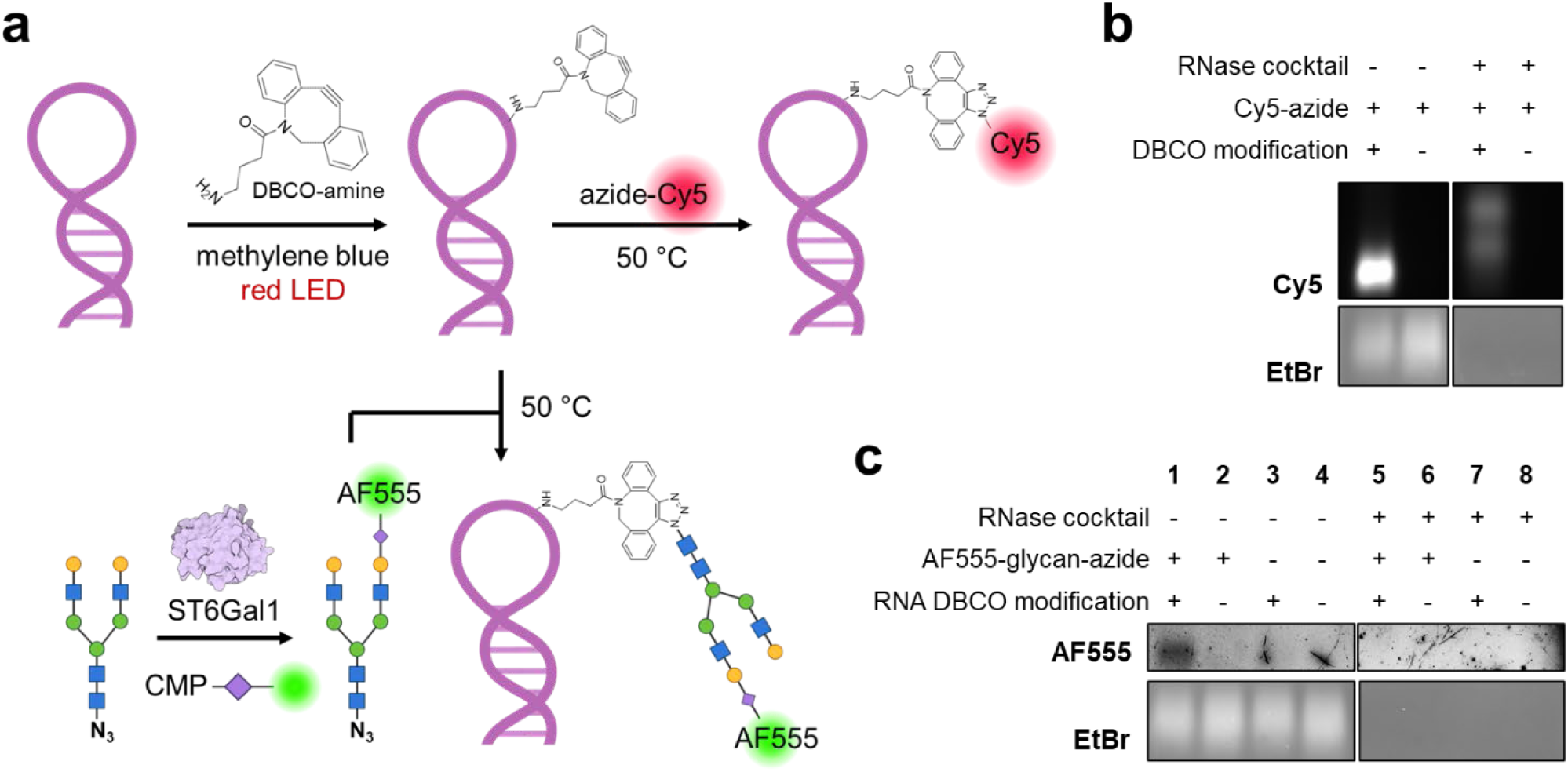
Chemically prepared neo-glycoRNA has different properties from N-glycoconjugates. **a.** Schematics for preparing DBCO-functionalized RNA and N-glycan conjugated neo-glycoRNA. Abbreviations: DBCO, dibenzocyclooctyne; LED, light-emitting diode; ST6Gal1, beta-galactoside alpha-2,6-sialyltransferase 1; CMP, cytidine monophosphate; AF555, AlexaFluor 555. **b.** 1% denaturing agarose gel electrophoresis and in-gel fluorescence of Cy5-clicked, DBCO-functionalized RNA. **c.** 1% denaturing agarose gel electrophoresis and in-gel fluorescence of neo-glycoRNA and RNase-treated samples. The AF555 signal has a highly similar mobility to EtBr signals.

To confirm the photochemical incorporation of DBCO, we also included a positive control, in which an azide-funcitonalized fluorophore (Cy5) was directly clicked with DBCO-conjugated RNA (**Fig.3a**). The chemically modified RNA samples were subjected to agarose gel electrophoresis after the purification. The strong Cy5 signal demonstrated the incorporation of DBCO into the RNA photochemically (**Fig.3b**). Little, if any, non-specific interaction between the dye and the RNA was observed. In stark contrast to glycoRNA or the N-glycoconjugates, the neo-glycoRNA had a similar apparent molecular weight in 1% agarose gel to the unmodified counterparts (**Fig.3c**, Lane 1), as the fluorescent band migrated to the similar position to the EtBr-stained bands. The result indicates the attachment of a sialylated N-glycan to RNA did not substantially alter the apparent molecular weight. Furthermore, the fluorescent band was lost upon RNase treatment right before sample loading into gels (Lane 5), suggesting N-glycans do not confer any RNase resistance to modified RNA. The results on the neo-glycoRNA further support the N-glycoconjugate does not have the chemical nature of RNA. The results also suggest additional structural factors may account for the different electrophoretic properties between neo-glycoRNA and real glycoRNA^28^.

### More alcohol enhances the silica adsorption of RNA-associated N-glycoconjugate

We sought to understand why the N-glycoconjugate was poorly recovered using silica column purification, especially when RNA is absent. We reason that the recovery must be governed by the working principle of silica-based solid phase: A more polar solution elutes the adsorbate faster. In the case of the N-glycoconjugate, 50% (v/v) ethanol in the sample loading buffer (suggested by vendors) is too polar for the N-glycoconjugate to be adsorbed, and likely has eluted most of it directly from the column before the subsequent washing steps (by buffers with 80% ethanol).

Based on this, we varied the percentage of ethanol or isopropanol from 20% to 80% in the sample loading buffer when purifying RNase-treated and untreated samples using the silica column (**Fig. 4a**). We found that the N-glycoconjugate was not captured at ethanol concentrations below 40%, regardless of RNase treatment. In line with our previous observations, at 50% ethanol concentration, the glycoconjugate was recovered in untreated conditions but not in the RNase-treated condition (**Fig. 4b** and **c** and **Extended Data Fig. 4**). When the percentage of ethanol increased, the glycoconjugate was recovered more efficiently (**Fig. 4b** and **c** and **Extended Data Fig. 4**). At around 60% ethanol concentrations and above, the glycan signal became visible even in the RNase-treated condition (**Fig. 4b** and **c** and **Extended Data Fig. 4b**). Similar results were obtained when ethanol was replaced with isopropanol (**Fig. 4b**). Our results on the correlation between the polarity of the loading buffer and the recovery efficiency of the N-glycoconjugate pointed to a co-adsorption or co-precipitant effect of RNA on the former and further strengthened our claims that the N-glycoconjugate had been physically depleted over the silica column in the absence of RNA, but not digested by RNase.

**Figure 4.**
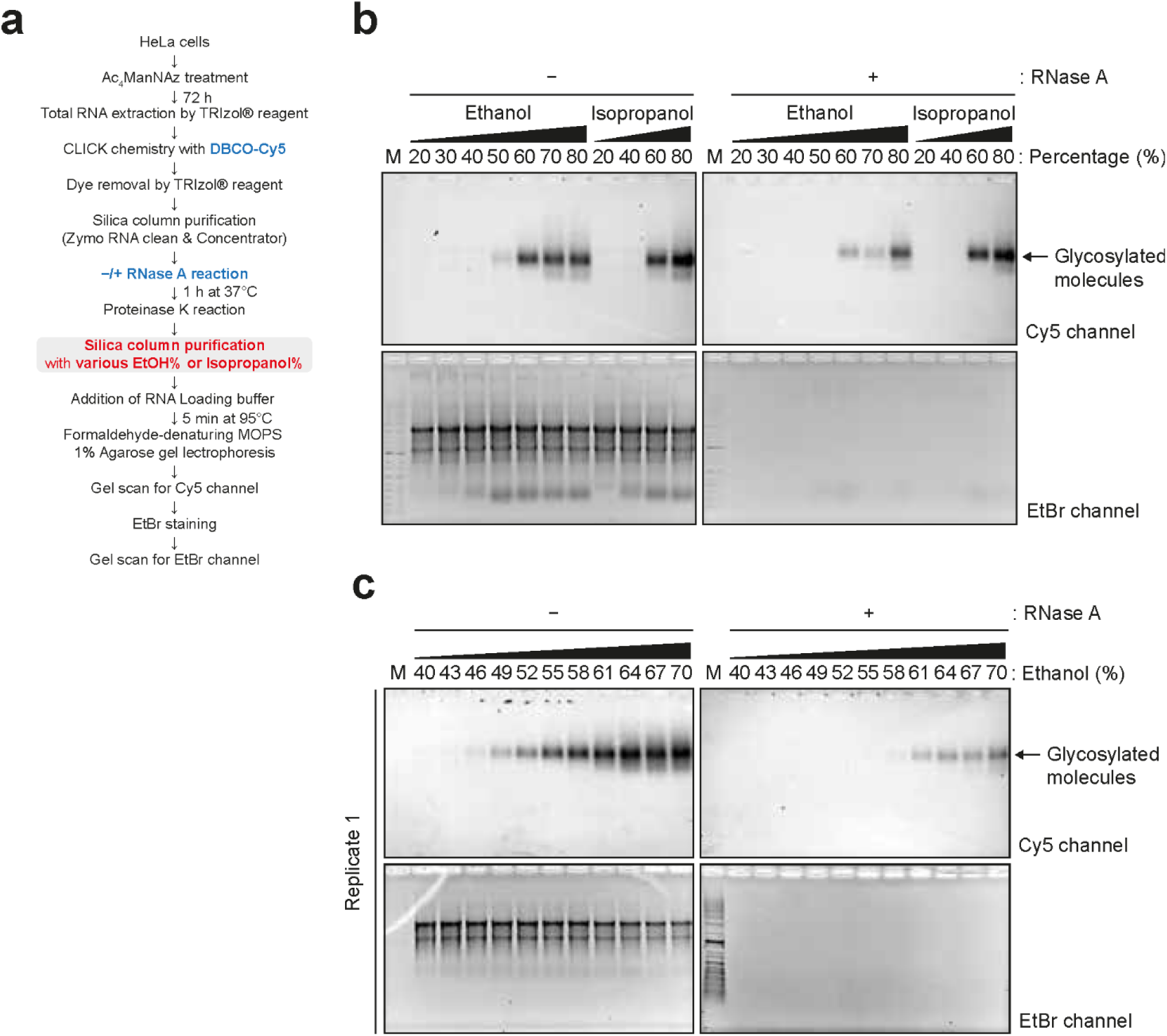
Ethanol percentage and RNA existence are critical for glycosylated molecule binding onto silica column. **a.** Schematic of glycan visualization protocol for various ethanol or isopropanol percentage in silica column binding solutions. **b.** Glycan detection in 20–80% ranges of ethanol or isopropanol by 10% increments. **c.** Glycan detection in 40–70% ranges of ethanol by 3% increments.

### Exogenous RNA facilitates the adsorption of the N-glycoconjugate on silica columns

Our observation indicates that the N-glycoconjugate in RNA preparations are not digested by RNase but did become less adsorbed on silica columns at around 50% ethanol concentration in the loading buffer when RNA had been degraded. This led us to the hypothesis that the total RNA, regardless of sources, has an enhancing effect on the silica adsorption of the N-glycoconjugate. On this basis, we should expect a heightened recovery of the N-glycoconjuagte over silica column when exogenous RNA is added back in the loading buffer. We thus performed a rescue experiment by adding total RNA extracted from unlabeled HeLa cells (i.e., not exposed to Ac_4_ManNAz) to the RNase treated sample (**Fig. 5a**). To ensure newly added RNA was not degraded by residual RNase activity, we removed RNase thoroughly by treating the samples with proteinase K followed by TRIzol extraction before the addition of new total RNAs. Strikingly, exogenously added, unlabeled total RNA effectively reduced the required minimum ethanol percentage for binding of the N-glycoconjugate to silica columns (**Fig. 5b**). Adding only one-tenth of the amount of RNA typically present in our RNA preparation was sufficient to fully restore recovery (**Fig. 4b**). Similarly, unlabeled RNA from a different cell line (K562) also enhanced binding efficiency of the N-glycoconjugate (**Extended Data Fig. 5a** and **b**).

**Figure 5.**
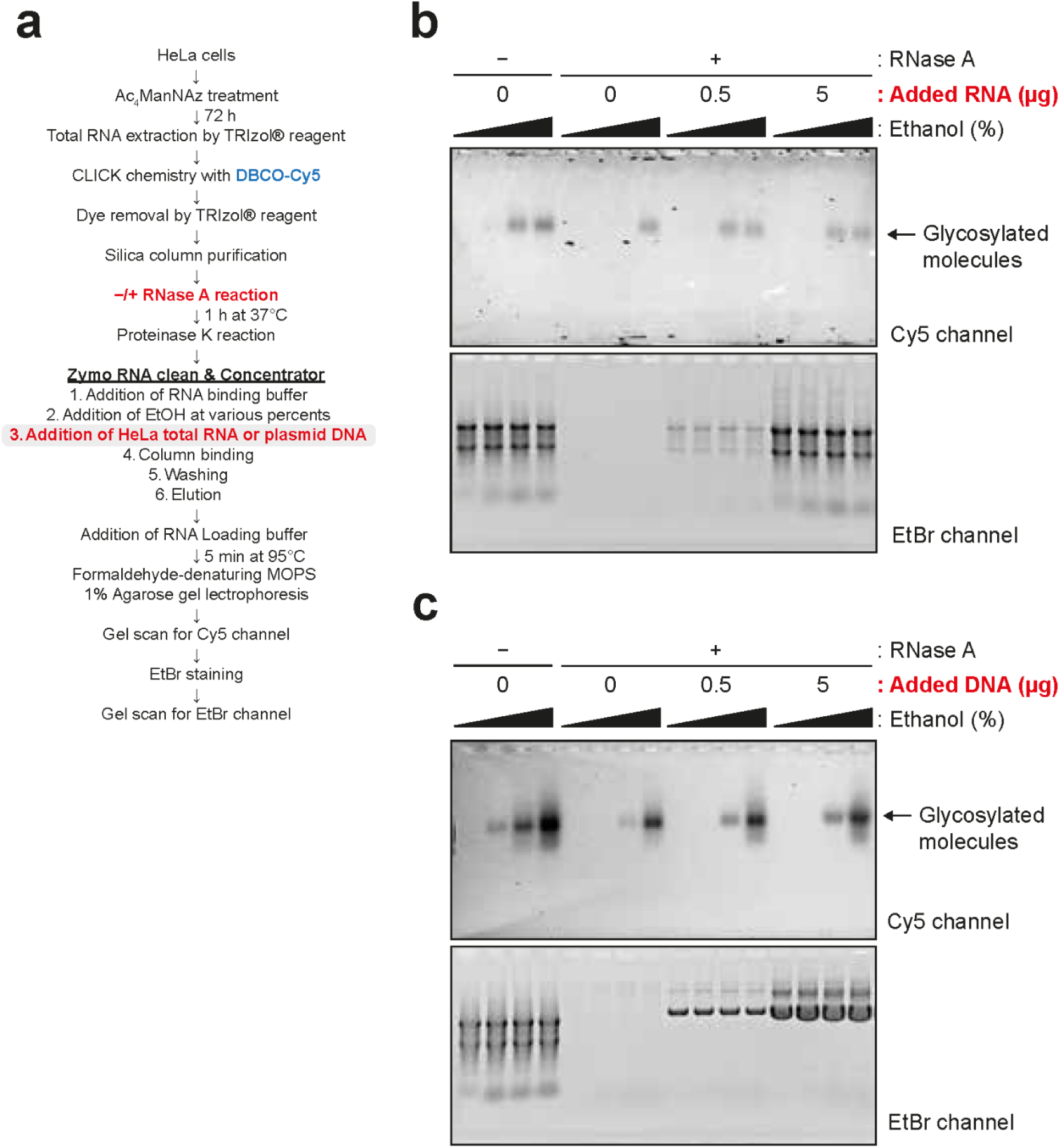
RNA is the co-binder of glycosylated molecules during silica column purification. **a.** Schematic of glycan visualization protocol for RNA or DNA addition in silica column binding solutions. 5 µg of total RNA extracted from Ac_4_ManNAz-treated HeLa cells were subjected to click chemistry per sample. **b.** Glycan detection without or with total RNA addition in RNA-depleted samples. The amounts of added total RNAs extracted from DMSO-treated HeLa cells are indicated. The range of the ethanol concentrations is 40–70%. **c.** Glycan detection without or with plasmid DNA addition in RNA-depleted samples. The amounts of added plasmid DNA are indicated. The range of the ethanol concentration is 40–70%.

Moreover, partially fragmented RNA was as sufficient in co-precipitating the N-glycoconjuagte (**Extended Data Fig. 5c** and **d**). However, more intense RNA fragmentation, which resulted in almost complete RNA degradation, did not have the enhancing effect on N-glycoconjugate recovery, an observation akin to RNase treatment. Adding exogenous total RNA to the RNA-removed (by non-enzymatic fragmentation) samples also rescued the binding of the N-glycoconjugate to silica matrix (**Extended Data Fig. 5e**). Interestingly, the addition of plasmid DNA had little effect on the minimum ethanol percentage required for binding of the N-glycoconjugate to silica (**Fig. 5c**). These results suggest RNA, but not double-stranded DNA, may interact with the N-glycoconjugate, leading to their efficient co-isolation in silica-column based extraction methods. Our rescue experiment also demonstrates that the N-glycoconjugate is unlikely covalently attached to RNA, despite its persistence throughout RNA preparations.

## Discussion

The recent description of N-glycosylated small RNA molecules has shaken our view on the principles of glycosylation and expanded the repertoire of post-transcriptional RNA modifications ^1^. Despite at its infancy, the glycoRNA field has recently seen considerable developments regarding its biological functions in cancer and innate immunity^4,5^, as well as the indication of its chemical nature^23^. In all studies, biochemically characterizing glycoRNA using electrophoresis has been essential for supporting the conclusions in the published work. Our findings serve as important heads-up for the fast-developing field, describing that a non-RNA N-glycoconjugate as an independent, separate molecular entity with broadly similar properties to glycoRNA may confound the biochemical analysis of the latter. It is also crucial to note that in our work, we initiated our investigation employing a procedure independent of the one in prior publications, attempting to maximize the recovery of glycoRNA, and presuming that independent procedures should serve to strengthen a biological observation. We compared our procedure with the reported workflow (referred to as “late-click” procedure in this manuscript). Although we have had consistent observations across both procedures, the higher yield of total RNA together with any associated molecular entities justified following up on our own procedure.

The non-RNA N-glycoconjugates persist in RNA preparations from mammalian cells even after multiple clean-ups and are resistant to digestion by multiple enzymes, including RNases, DNases and proteinase K. The N-glycoconjugates also remain with RNA during phenol-chloroform phase separation and ethanol precipitation-based purification. The non-RNA nature of the N-glycoconjugates was further indicated by their drastic difference from the chemically prepared neo-glycoRNA, in which the attachment of glycans did not confer RNA the protection from degrading enzymes. However, these persisting N-glycoconjugates did become depleted, but not digested, only when using silica columns to purify the RNase-treated samples. Given that assays on glycoRNA also routinely employ silica columns for the final clean-up before electrophoresis, it is thus difficult to distinguish the two scenarios when the loss of the glycan-labeled bands is observed: Should the loss of signals be interpreted as 1) RNase digestion of glycoRNA or 2) the physical depletion of N-glycoconjugates upon RNA removal?

To address this question, we propose a simple checkpoint experiment for the relevant fields, as was performed in our study. With this, one would be able to distinguish if the desired molecule is being studied. We suggest treating the purified, glycan-labeled post-click RNA samples with RNases for an extended period, and then directly load the mixture into the gel for electrophoresis, without using a column to clean up the sample, regardless of whether the sample preparation follows an early-click or a late-click procedure as reported. With a transfer to membrane or not, the band at large molecular weight should disappear for glycoRNA. If it does not, one should be alerted that the non-RNA N-glycoconjugates have been largely co-isolated.

Our observation prompted us to investigate the mechanism of the association of the N-glycoconjugates during silica-column purification. Firstly, with varying ethanol concentration or switching to isopropanol during sample loading, we observed that the N-glycoconjugate adsorption follow the general rules for silica-based systems: the more apolar content (i.e. ethanol or isopropanol) in the loading buffer, the better the molecules are retained on the silica resin. Although 60% or higher ethanol, or isopropanol, in the loading buffer deviates from the vendor-suggested alcohol content for RNA sample loading, it did substantially improve the recovery of the N-glycoconjugates over a silica column, especially compared to that when RNA had been removed before sample loading. Notably, glycoRNA has been shown to exhibit similar adsorption properties on silica^23^. Secondly, the N-glycoconjugate is not covalently attached to RNA. However, exogenously added RNA regardless of sources of extraction can enhance the adsorption of the N-glycoconjugates to silica resin, indicating that RNA acts as a co-precipitant for the N-glycoconjugates.

While the chemical nature of the newly characterized N-glycoconjugates in this work remain to be elucidated, we propose N-glycosylated peptides as a candidate which warrant follow-up confirmative studies. N-glycans themselves may contain negatively charged moieties, such as sialic acid, phosphate, or sulfate groups ^29–34^, which may contribute to their mobility towards the positive electrode during gel electrophoresis and efficient transfer to a positively charged nylon membrane. Thus, potential candidates of these molecules may be N-glycosylated oligopeptide products degraded from glycoproteins, which cannot be further cleaved by proteinase K. The possible peptidic nature of glycoRNA-associated glycosylated molecules is supported by the cleavage of an N-glycan from asparagine residues by PNGase F, which requires at least a tripeptide-containing substrate ^35^. Interestingly, a highly hydrophilic oligopeptide containing a sialylated complex-type N-glycan linked to a hexapeptide can be isolated from chicken egg yolk in considerable quantity and high homogeneity ^25^. It is thus intriguing to ask if similar molecules also exist in mammalian cells.

Of general importance, our findings demonstrate that even the gold standard RNA purification methods are susceptible to seemingly inert molecules such as glycans which are not easily detected by conventional means. It should be brought to the attention that co-isolation of other biomolecules with extracted nucleic acids are not uncommon. For example, anti-coagulant heparin often contaminates purified DNA or RNA samples from blood collection and plasma processing procedures, and such contamination can complicate reverse transcription and PCR analysis ^14,15^. It is currently unclear how glycosylated molecules may have affected and will affect studies that have relied on conventional RNA isolation methods. Our work prompts the development of more reliable RNA purification and post-transcriptional modification methods and will serve as a catalyst for further investigation into a potentially novel biomolecule.

## METHODS

### Key resources table

| Reagent or resource | Source | Identifier |
| --- | --- | --- |
| **Chemicals, peptides, and recombinant proteins** |  |  |
| Dulbecco's Modified Eagle's Medium (DMEM), high glucose with L-glutamine, sodium pyruvate and sodium bicarbonate | Welgene | Cat# LM001-17 |
| Dulbecco's Modified Eagle's Medium (DMEM) | Gibco | Cat# 41965-039 |
| Fetal bovine serum | Welgene | Cat# S001-01 |
| Fetal bovine serum | Sigma-Aldrich | Cat# F7524 |
| HyClone Characterized Fetal Bovine Serum | Cytiva | Cat# SH30071.03HI |
| Penicillin/streptomycin | Sigma-Aldrich | Cat# P4458 |
| Dulbecco's modified phosphate buffered saline (D-PBS), without calcium chloride and magnesium chloride | Welgene | Cat# LB001-02 |
| Phosphate buffered saline (PBS) | In-house preparation |  |
| Dimethyl sulfoxide (DMSO) | Sigma-Aldrich | Cat# 276855 |
| Dimethyl sulfoxide (DMSO) | Merck | Cat# 1.0295.1000 |
| N-azidoacetylmannosamine-tetraacylated (Ac<sub>4</sub>ManNAz) | Sigma-Aldrich | Cat# 900917-50MG |
| N-azidoacetylmannosamine-tetraacylated (Ac<sub>4</sub>ManNAz) | Custom synthesis by Synvenio |  |
| TRIzol™ Reagent | Thermo Fisher Scientific | Cat# 15596018 |
| TRI Reagent™ Solution | Thermo Fisher Scientific | Cat# AM9738 |
| TRIzol™ LS Reagent | Thermo Fisher Scientific | Cat# 10296028 |
| Chloroform | Sigma-Aldrich | Cat# C2432 |
| TURBO™ DNase (2 U/μL) | Thermo Fisher Scientific | Cat# AM2238 |
| DNase I (RNase free; 2U/μl) | Thermo Fisher Scientific | Cat# AM2222 |
| RNase A (DNase and protease-free, 10 mg/mL) | Thermo Fisher Scientific | Cat# EN0531 |
| PureLink™ RNase A (20 mg/mL) | Thermo Fisher Scientific | Cat# 12091021 |
| Rapid PNGase F | New England Biolabs | Cat# P0710S |
| α2-3,6,8,9 Neuraminidase A | New England Biolabs | Cat# P0722S |
| Proteinase K, Recombinant, PCR grade | Roche | Cat# 3115879001 |
| Proteinase K | Thermo Fischer Scientific | Cat# AM2548 |
| RNase T1 | Thermo Fischer Scientific | Cat# AM2283 |
| RNase cocktail | Thermo Fischer Scientific | Cat# AM2286 |
| Benzonase® Nuclease | Merck | Cat# E1014-25KU |
| RNase H | New England Biolabs | Cat# M0297L |
| Nuclease P1 | New England Biolabs | Cat# M0660S |
| UltraPure™ Formamide | Thermo Fisher Scientific | Cat# 15515026 |
| UltraPure™ 0.5M EDTA, pH 8.0 | Thermo Fisher Scientific | Cat# 15575020 |
| DEPC-Treated H<sub>2</sub>O | Thermo Fisher Scientific | Cat# AM9920 |
| Isopropanol, Optima LC/MS Grade | Fisher Scientific | Cat# A461-500 |
| Ethyl alcohol, Pure | Sigma-Aldrich | Cat# E7023-1L |
| dibenzocyclooctyne-Cy5 (DBCO-Cy5) | Sigma-Aldrich | Cat# 777374-5MG |
| dibenzocyclooctyne-PEG4-biotin (DBCO-biotin) | Sigma-Aldrich | Cat# 760749-5MG |
| Linear acrylamide | Thermo Fisher Scientific | Cat# AM9520 |
| SeaKem® LE Agarose | Lonza | Cat# 50004 |
| NorthernMax® Denaturing Gel Buffer (10X) | Thermo Fisher Scientific | Cat# AM8676 |
| NorthernMax® 10X Running Buffer | Thermo Fisher Scientific | Cat# AM8671 |
| NorthernMax® Transfer Buffer | Thermo Fisher Scientific | Cat# AM8672 |
| Odyssey Blocking Buffer (PBS) | Li-Cor Biosciences | Cat# 927-40000 |
| IRDye 800CW Streptavidin | Li-Cor Biosciences | Cat# 926-32230 |
| MOPS | Sigma-Aldrich | Cat# M1254 |
| PBS Tablets | Gibco | Cat# 18912-014 |
| TWEEN® 20 | Sigma-Aldrich | Cat# P7949 |
| Sodium Acetate·3H<sub>2</sub>O | Merck | Cat# 1.06265.1000 |
| EDTA | Sigma-Aldrich | Cat# EDS |
| 37% Formaldehyde solution | Merck | Cat#1.04003.1000 |
| UltraPure™ Ethidium Bromide, 10 mg/mL | Thermo Fisher Scientific | Cat# 15585011 |
| Zeta-Probe® GT Membrane | Bio-Rad | Cat# 1620194 |
| BrightStar™-Plus Positively Charged Nylon Membrane | Thermo Fisher Scientific | Cat# AM10102 |
| Hybond-C nitrocellulose membrane | Cytiva | Cat# RPN303C |
| Amersham™ Protran® nitrocellulose membrane | Merck | Cat# GE10600001 |
| Ambion® 10X RNA Fragmentation Reagent | Thermo Fisher Scientific | Cat# AM8740 |
| **Critical commercial assays** |  |  |
| NorthernMax™ Kit | Thermo Fisher Scientific | Cat# AM1940 |
| RNA Clean and Concentrator 5 | Zymo Research | Cat# R1013 |
| **Experimental models: cell lines** |  |  |
| HeLa | ATCC | Cat# ATCC-CCL-2 |
| K562 | ATCC | Cat# ATCC-CCL-24 |

### Lead contact

Further information and requests for resources and reagents should be directed to and will be fulfilled by the Lead Contact, Sungchul Kim.

### Materials availability

All unique/stable reagents generated in this study are available from the Lead Contact with a completed Materials Transfer Agreement.

## Experimental Model and Subject Details

### Mammalian cell culture

HeLa cells were cultured at 37°C and 5% CO_2_ in Dulbecco’s Modified Eagle’s Medium (DMEM, Welgene, Fig. 1b,c, 2b–f, 3b,c and 4b,c and Extended Data Fig. 1b,c, 2a–d, 3a and 4d,e; Gibco, Extended Data Fig. 2b) media supplemented with 10% fetal bovine serum (FBS) (Welgene, Fig. 1b,c, 2b–f 3b,c and 4b,c and Extended Data Fig. 1b,c, 2a,c,d, 3a and 4d,e; Sigma-Aldrich, Extended Data Fig. 2b), and also supplemented with penicillin/streptomycin (P/S) (Sigma-Aldrich, Extended Data Fig. 2b). Cells were maintained in 100-mm cell culture dishes (Fig. 1b,c, 2b–f 3b,c and 4b,c and Extended Data Fig. 1b,c, 2a,c,d, 3a and 4d,e) with 10 mL of culture media or, in T75 flasks (Extended Data Fig. 2b). When reaching confluency, cells were split for sub-culturing. K562 cells (Extended Data Fig. 4b) were cultured at 37°C and 5% CO_2_ in RPMI medium 1640 supplemented with 10% FBS (Cytiva) and P/S (Sigma-Aldrich) with shaking at 120 rpm.

## Method details

### Labeling with metabolic chemical reporter

Stocks of 500mM N-azidoacetylmannosamine-tetraacylated (Ac_4_ManNAz) (Sigma-Aldrich, Fig. 1b,c, 2b–f, 4b,c and 5b,c and Extended Data Fig. 1b,c, 2b–d, 4a and 5b,d,e) were prepared in sterile dimethyl sulfoxide (DMSO) (Sigma-Aldrich). For metabolic labeling, we treated Ac_4_ManNAz at a final concentration of 100 µM in fresh DMEM supplemented with 10% FBS. After 72 h incubation at 37°C and 5% CO_2_, cells were washed with Dulbecco’s Phosphate-Buffered Saline (D-PBS) (Welgene, Fig. 1b,c, 2b–f, 4b,c and 5b,c and Extended Data Fig. 1b,c, 2b–d, 4a and 5b,d,e) twice and stored in -80°C until total RNA extraction. For experiments shown in Extended Data Fig. 4b, the conditions for metabolic labeling were the same, except for that P/S were added in the media.

For experiments shown in Extended Data Fig. 2b, of 50 mM stocks of Ac_4_ManNAz (Custom synthesis by Synvenio) were prepared in sterile DMSO (Merck). The metabolic labeling was done with 100 µM Ac_4_ManNAz. In the DMEM (Gibco) supplemented with 10% FBS (Sigma-Aldrich) and 1% P/S. After 48 h at 37 °C and 5% CO_2_, cells were washed with PBS (in-house preparation) and then followed by total RNA extraction.

### Total RNA extraction with TRIzol™ Reagent

For experiments shown in Fig. 1b,c, 2b–f, 4b,c and 5b and 5c and Extended Data Fig. 1b,c, 2b–d, 4a and 5b,d,e, 1 mL of TRIzol™ reagent (Thermo Fisher Scientific) was added directly onto a washed cell culture dish. Dishes were rocked thoroughly for 10 min at room temperature to lyse and denature all cells. For extracting total RNA from K562 cells (Extended Data Fig. 4b), the cell pellets washed twice with D-PBS were lysed in TRIzol™ reagent (Thermo Fisher Scientific) with 1 mL for ∼10^7^ cells). Homogenized TRIzol-cell mixtures were scrapped with cell scrapper and then transferred into nuclease-free sterile 1.7 mL microtubes. The tubes were vortexed at least for 5 min until complete homogenization for further denaturation of the intermolecular non-covalent interactions. Phase separation was initiated by adding 200 µL (0.2× volumes) of 100% chloroform (Sigma-Aldrich) into 1-mL TRIzol-dissolved cell mixture, and then vortexed thoroughly for complete homogenization. The homogenates were centrifuged at 16,000×g for 10 min at 4°C. The upper (aqueous) phase was carefully removed, transferred into a nuclease-free sterile 1.7 mL, and then mixed with equal volume of 100% isopropanol (Fisher Scientific) by vortexing. The mixture was centrifuged at 16,000×g for 30 min at 4°C, and the supernatant was carefully discarded. The pellet was washed with 1 mL of ice-cold 75% ethanol (Sigma-Aldrich) twice, and then dried completely. The RNA pellet was dissolved with Milli-Q^®^ (Thermo Fisher Scientific, Fig. 1b,c, 2b–f, 3b,c and 4b,c and Extended Data Fig. 1b,c, 2b–d, 4a and 5b,d,e) H_2_O or DEPC-treated H_2_O (Thermo Fisher Scientific, Extended Data Fig. 5b), and the concentration was measured using a Nanodrop™ 2000 UV/Vis spectrophotometer (Thermo Fisher Scientific).

For experiments shown in Extended Data Fig. 2b, 6 mL of TRI Reagent Solution (Thermo Fisher Scientific) was added directly to the washed T75 cell culture flask. Homogenates were vortexed for 1 min at RT followed by incubating the samples 1 min at 37°C. 0.2× volumes of chloroform were added, and phase separation was performed at 16,000×g for 10 min at RT. After adding equal volume of isopropanol, mixtures were incubated for 10 min at 4°C. RNA was precipitated at 16,000×g for 10 min at 4°C, washed twice with 75% ethanol and dissolved in nuclease-free H_2_O. To obtain highly pure RNA preparations, The isolated RNA was re-purified by adding 1 ml of TRI Reagent Solution (Thermo Fisher Scientific) and repeating the isolation procedure described above.

### Copper-free click chemistry and removal of free ligands

Strain-promoted alkyne-azide cycloadditions (SPAAC) was performed to probe for azide-containing sialo-conjugates in RNA samples using dibenzocyclooctyne-conjugated cyanine 5 (DBCO-Cy5) (Sigma-Aldrich) dyes or DBCO-biotin (Sigma-Aldrich) as the alkyne-azide cycloaddition. Stocks of 10 mM DBCO-Cy5 or DBCO-biotin were made by dissolving 1 mg of lyophilized DBCO-Cy5 in 82.6 µL or 5 mg of DBCO-biotin in 666.7 µL of DMSO, respectively. 9 µL (typically ∼50 µg) of RNA dissolved in H_2_O were mixed at first with 10 µL of home-made 2× dye-free RNA loading buffer (df-RLB, 95% formamide, 25 mM EDTA, pH8.0) and added with 1 µL of 10 mM (for final 500 µM) DBCO-Cy5 or DBCO-biotin in a microtube. Samples were incubated at 55°C for 10 min. The reaction was stopped by adding 1 mL of TRIzol™ reagent (Thermo Fisher Scientific) and 200 µL of chloroform (Sigma-Aldrich). Alternatively, for experiments performed in Extended Data Fig. 2b, dye removal was achieved by adding 80 µL of DEPC-treated H_2_O, 300µl of TRIzol LS Reagent (Thermo Fisher Scientific) and 80 µL of Chloroform (Merck). Samples were centrifuged at 16,000×g for 10 min at 4°C or, for experiments shown in Extended Data Fig. 2b, at 16,000×g for 5 min at room temperature. The upper (aqueous) phase was carefully removed, transferred into a nuclease-free sterile tube. For Fig. 1b,c, 2b–f, 4b,c and 5b,c and Extended Data Fig. 1b,c, 2a–d, 4a and 5b,d,e, samples were mixed with equal volume of 100% isopropanol by vortexing, subsequently centrifuged at 16,000×g for 30 min at 4°C, and the supernatant was carefully discarded. The pellet was washed with 1 mL of ice-cold 75% ethanol twice, and then dried completely. The RNA pellet was dissolved with Milli-Q^®^ H_2_O (Fig. 1b,c, 2b–f, 4b,c and 5b,c and Extended Data Fig. 1b,c, 2a–d, 4a and 5d,e) or DEPC-treated H_2_O (Extended Data Fig. 5b), and the concentration was measured using the UV/Vis spectrophotometer.

### Enzymatic reactions

Typically, enzymatic reactions were performed with 10 µg (Fig. 2b–f, 4b,c and 5b,c and Extended Data Fig. 1c, 2a– d, 4a and 5b) or 12 µg (Extended Data Fig. 2b) of labeled RNA at 37°C. To digest RNA, 0.5 µL of RNase A (DNase and protease-free, 10 mg/mL) (Thermo Fisher Scientific, Fig. 2b–f, 3b,c and 4b,c Extended Data Fig. 1c, 2a–d, 3a and 4b; PureLink™ RNase A (20 mg/mL) (Thermo Fisher Scientific, Extended Data Fig. 4b), or 1 µL of RNase T1 (Thermo Fisher Scientific, Fig. 2f and Extended Data Fig. 2c,d), or 1 µL of RNase cocktail (500 U of RNase A and 20,000 U of RNase T1 per mL) (Thermo Fisher Scientific, Fig. 2f and Extended Data Fig. 2c,d) were treated with 1.5 µL of 10× TURBO DNase buffer (Thermo Fisher Scientific) in the sample by adjusting with Milli-Q H_2_O to total 15 µL. To degrade DNA, 0.5 µL of TURBO DNase (2,000 U/mL) (Thermo Fisher Scientific, Fig. 2b–f and Extended Data Fig. 1c and 2a,c,d) or DNase I (2,000 U/mL) (Thermo Fisher Scientific, Extended Data Fig. 2b) were treated with 1.5 µL of 10× DNase buffer in the sample by adjusting with Milli-Q H_2_O to total 15 µL. To digest both RNA and DNA, 1 µL of benzonase (250,000 U/mL) (Merck, Fig. 2f and Extended Data Fig. 2c,d) or 1 µL of nuclease P1 (100,000 U/mL) (New England Biolabs, Fig. 2f and Extended Data Fig. 2c,d) were treated with 1.5 of 10× TURBO DNase buffer or nuclease P1 buffer in the sample by adjusting with Milli-Q H_2_O to total 15 µL. To digest RNA in DNA/RNA hybrids, 1 µL of RNase H (5,000 U/mL) (New England Biolabs, Fig. 2f and Extended Data Fig. 2c,d) were treated with 1.5 of 10× RNase H buffer in the sample by adjusting with Milli-Q H_2_O to total 15 µL. To digest N-glycans, 0.5 µL of Rapid PNGase F (New England Biolabs, Fig. 2d and Extended Data Fig. 2a) were added with 1.5 µL of 10× PNGase F buffer (New England Biolabs) in the sample by adjusting with Milli-Q H_2_O to total 15 µL. To cut sialic acid moieties, 0.5 of α2-3,6,8,9 Neuraminidase A (New England Biolabs, Fig. 2d and Extended Data Fig. 2a,b) were added with 1.5 µL of 10× GlycoBuffer 1 (New England Biolabs) in the sample by adjusting with Milli-Q H_2_O to total 15 µL. To digest proteins, 1 µL of proteinase K (PK, Roche, 20 mg/mL dissolved in Milli-Q H_2_O, Fig. 1b,c, 2d–f, 4b,c and 5b,c and Extended Data Fig. 1b,c, 2a–d, 4a and 5b; Thermo Fischer Scientific, 20 mg/mL, Extended Data Fig. 2b) was added either with 1.5 µL of 10× TURBO DNase buffer in the RNA sample by adjusting with Milli-Q H_2_O to total 15 µL or directly in the precedent enzymatic reactant. The incubation was done for 30 min or 60 min in cases, but all the results always exhibited complete protein digestion.

### RNA fragmentation

DBCO-Cy5-labeled RNA was fragmented using Ambion^®^ 10X RNA Fragmentation Reagent (Thermo Fisher Scientific). Samples were incubated in 1× RNA Fragmentation Reagent at 75°C for 15 min for mild reaction (Extended Data Fig. 5d) and at 95°C for 2 h for complete fragmentation (Extended Data Fig. 5e). Fragmented RNA samples were immediately mixed with 2× volumes of RBB and various volumes of 100% ethanol for each sample for the final 40%, 50%, 60% and 70% ethanol as indicated in Extended Data Fig. 4d,e.

### RNA clean-up by acidic phenol-chloroform extraction

For experiments in Fig. 2b,d,f and Extended Data Fig. 2a,c,d, enzymatically digested samples were mixed thoroughly with 1 mL of TRIzol reagent and 200 µL of 100% chloroform (Sigma-Aldrich) for 10 min at room temperature. The homogenates were centrifuged at 16,000×g for 10 min at 4°C. The upper phase was carefully removed, transferred into a nuclease-free sterile 1.7-mL tube, and then mixed with 1 µL of linear acrylamide (Thermo Fisher Scientific) as a co-precipitant and equal volume of 100% isopropanol by vortexing, subsequently centrifuged at 16,000×g for 30 min at 4°C, and the supernatant was carefully discarded. The pellet was washed with 1 mL of ice-cold 75% ethanol twice, and then dried completely. The pellet was dissolved with Milli-Q^®^ H_2_O. For experiments in Extended Data Fig. 2b, 20 µg linear acrylamide and TRI Reagent Solution (Thermo Fisher Scientific) were used.

### RNA clean-up and size fractionation by silica-based column purification

16 µL of PK-digested samples were mixed with 34 µL of Milli-Q H_2_O to total 50 µL. 100 µL of RBB and 150 µL of 100% ethanol (Final 50% ethanol) were added by reverse pipetting and vortexed thoroughly. For experiments shown in Extended Data Fig. 2b, the final percentage of EtOH was 60%. The mixtures were transferred into the Zymo-SpinTM IC Column in a 2 mL of collection tube. The columns were centrifuged at 16,000×g for 30 sec at room temperature and the flow-through was discarded. 400 µL of RNA Prep Buffer (RPB) (provided by RNA Clean & Concentrator-5, Zymo Research) were added into the column and then centrifuged at 16,000×g for 30 sec at room temperature followed by discarding the flow-through. 700 µL of RNA Wash Buffer (RWB) (provided by RNA Clean & Concentrator-5, Zymo Research) were added into the column and then centrifuged at 16,000×g for 30 sec at room temperature followed by discarding the flow-through. Add 400 µL of RWB were added into the column and then centrifuged at 16,000×g for 30 sec at room temperature followed by discarding the flow-through. Centrifuge at 16,000×g for 1 min at room temperature again to ensure complete removal of the RWB. The columns were carefully transferred into a new nuclease-free sterile 1.7-mL tube, and 15 µL of Milli-Q H_2_O or DEPC-treated H_2_O directly to the column matrix and incubate for 3 min. The elution was done by centrifuge at 16,000×g for 3 min at room temperature.

For size fractionation of small (smaller than about 200 nucleotides) versus large (larger than about 200 nucleotides) RNAs, samples mixed with equal volume of 50% RBB in 100% ethanol. The mixture was applied onto the Zymo-SpinTM IC Column and centrifuged at 16,000×g for 1 min at room temperature. Large RNAs bound in the column were purified as described above. Small RNAs in the flow-through were mixed with equal volume of 100% ethanol and then purified as described above.

### Photochemical labelling of fragmented RNA with amine-DBCO and click chemistry

The purified RNA sample was fragmented using NEBNext Magnesium RNA Fragmentation Module (NEB, E6150S) following protocols recommended by the vendor and purified using Zymo RCC-25. Amine-DBCO (CAS# 1255942-06-3) methylene blue (CAS# 61-73-4) were added to a solution of 50 µg RNA at a final concentration of 1 mM and 100 µM, respectively. The solution of was exposed to red light (50 W LED chip on board) on ice for 15 min with lid open. The resulted reaction mixture was then purified using TRIzol in combination with Zymo RCC-25 following the vendor-provided protocol. The purified DBCO-funcitonalized RNA (10 µg per reaction) was then clicked with either Cy5-azide (Jena Bioscience, CLK-1177) or the AlexaFluor-555 labelled sialoglycan-azide at 200 µM and 60 µM, respectively, at 50 °C for 10 min. The clicked RNA product was then purified with TRIzol in combination with Zymo RCC-10 following vendor’s protocol. The RNA samples were resolved with 1% denaturing agarose gel and fluorescently imaged under a GelDoc scanner, after which the gel was stained with EtBr and scanned again for total RNA.

To make AlexaFluor-555 labelled sialoglycan-azide, G2 glycoform N-glycan linked with an azido-asparagine (final concentration 120 µM) was mixed with 10 µg ST6Gal1 and 200 µM CMP-sialic acid pre-functionalized with AlexaFluor-555. The sialylation reaction was incubated at 37 °C for 48 h, and was used for the click chemistry with RNA directly without further purification.

### Denaturing gel electrophoresis, membrane transfer, and blotting

Typically, formaldehyde-denaturing 1% agarose gel was made by the following. 0.5 g of agarose powder (Lonza) were mixed in 45 mL of Milli-Q H_2_O in the flask by swirling but thoroughly. The flask was heated in the microwave oven until complete melting of the agarose. The flask was removed from the oven and then cooled to 55–60°C. 5 mL of 10× NorthernMax™ Denaturing Gel Buffer (Thermo Fisher Scientific) were added and mixed by swirling in the fume hood. The gel mixture was casted following the instruction provided by the casting apparatus. To prepare loading samples, samples were mixed with equal volume of 2× df-RLB, and then incubated at 95°C for 10 min. The gel was resolved in 1× NorthernMax™ Running Buffer (Thermo Fisher Scientific) at 100 V for 40–50 min. For visualization of the Cy5 fluorescence, the gel was rinsed briefly with Milli-Q H_2_O and scanned using the gel imaging system (Bio-Rad ChemiDoc XRS+) in the Cy5 filter channel. For ethidium bromide (EtBr) scanning, the Cy5 scanned gel was stained in the water-dissolved UltraPure™ EtBr (Thermo Fisher Scientific) solution for 30 min at room temperature by rocking. The gel was rinsed with Milli-Q H_2_O for 30 min at room temperature by rocking and then scanned in the gel imaging system.

For experiments shown in Extended Data Fig. 2b, 1 g of agarose powder (Roche) was dissolved in 72 mL of Milli-Q H_2_O. 10 mL of 10× MOPS buffer (200 mM MOPS, 50 mM sodium Acetate·3H_2_O, 10 mM EDTA, pH 7.0) and 18 mL 37% formaldehyde (Merck) were added and mixed. Purified, enzyme treated, RNA samples were incubated at 95 °C for 5 min followed by a quick transfer and 5 min incubation on ice before gel electrophoresis at 90V. Cy5 fluorescence was visualized using the Amersham Typhoon scanner (GE Healthcare).

For membrane transfer, the electrophoresed gel was scanned in the Cy5 channel and then immediately subjected to the transfer instead of EtBr staining, since the EtBr emission was strongly overlapped with Cy5 visualization during the downstream membrane scanning. NorthernMax™ Transfer Buffer (Thermo Fisher Scientific) was used following the manufacturer’s instruction for 2 h. For nylon membranes, Zeta-Probe^®^ GT (Bio-Rad) or BrightStar™-Plus (Thermo Fisher Scientific) membranes were used. For nitrocellulose membranes, Hybond-C (Cytiva) or Amersham™ Protran^®^ (Sigma-Aldrich) membranes were used. The transferred membranes were rinsed briefly with Milli-Q H_2_O and scanned immediately in the Cy5 channel using Bio-Rad ChemiDoc XRS+.

After transfer of the gel run with biotinylated samples, membranes were subjected to blocked with Odyssey Blocking Buffer, PBS (Li-Cor Biosciences) for 45 min at room temperature, by skipping EtBr staining and fluorescent imaging. After blocking, membranes were stained for 30 min at room temperature with streptavidin-conjugated IR800 (Li-Cor Biosciences), which was diluted to 1:5,000 in Odyssey Blocking Buffer. Excess IR800-streptavidin was washed from the membranes by four times with 0.1% TWEEN-20 (Sigma-Aldrich) in 1× PBS for 10 min/each at room temperature. Membranes were finally washed once with 1× PBS to remove residual TWEEN-20 before scanning. Fluorescent signals from membranes were scanned on Odyssey Li-Cor Sa scanner (Li-Cor Biosciences) with the software set to auto-detect the signal intensity for both 700 and 800 channels. After scanning, images were adjusted to appropriate contrasts with the Li-Cor software (when appropriate) in the 800 channel and exported.

## Acknowledgements

This work was supported by Young Scientist Fellowship program of the Institute for Basic Science from the Ministry of Science and ICT of Korea (IBS-R008-D1 to S.K.); by TU Delft-Leiden Health Initiative (V.R, C.J.); by a Dutch Research council (NWO) ENW-XS grant (OCENW.XS21.4.046 to P.M.); by ERC Consolidator grant of the European Research Council (819299 to C.J.); and by NWO VENI talent program (VI.Veni.222.272, Z.L.)

## Authors contributions

S.K. and Z.L. made the initial discovery. S.K., C.J. and Z.L. designed the study. S.K., Z.L., P.M., and C.J. jointly supervised the work. S.K., Y.C., K.J., A.P, and Z.L. performed the experiments. S.K., K.J., Z.L., P.M., and C.J co-wrote the manuscript with input from all authors.

---

**Extended Data Fig. 1.**
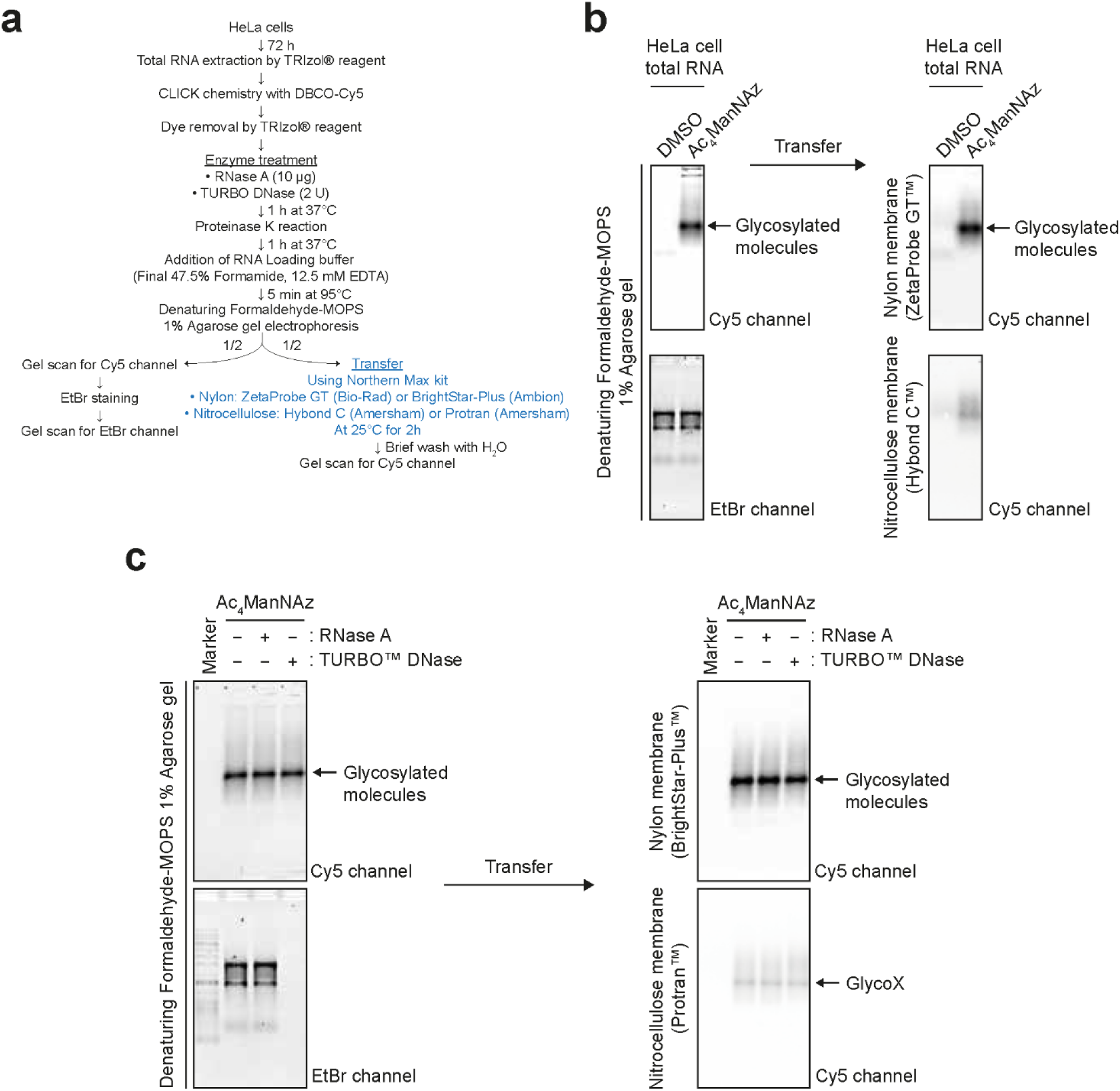
Membrane transfer assay for glycan detection, related to Figure 1. **a.** Schematics of the workflow. **b.** Glycan detection in formaldehyde-denaturing agarose gels and blotted membranes. For nylon membrane, ZetaProbe GT™ from Bio-Rad™ was used. For nitrocellulose membrane, Hybond C™ from AmershamTM was used. **c.** Glycan detection in gels and blotted membranes after RNase or DNase treatments. For nylon membrane, BrightStar-Plus™ from Invitrogen™ was used. For nitrocellulose membrane, Protran™ from Amersham™ was used.

**Extended Data Fig. 2.**
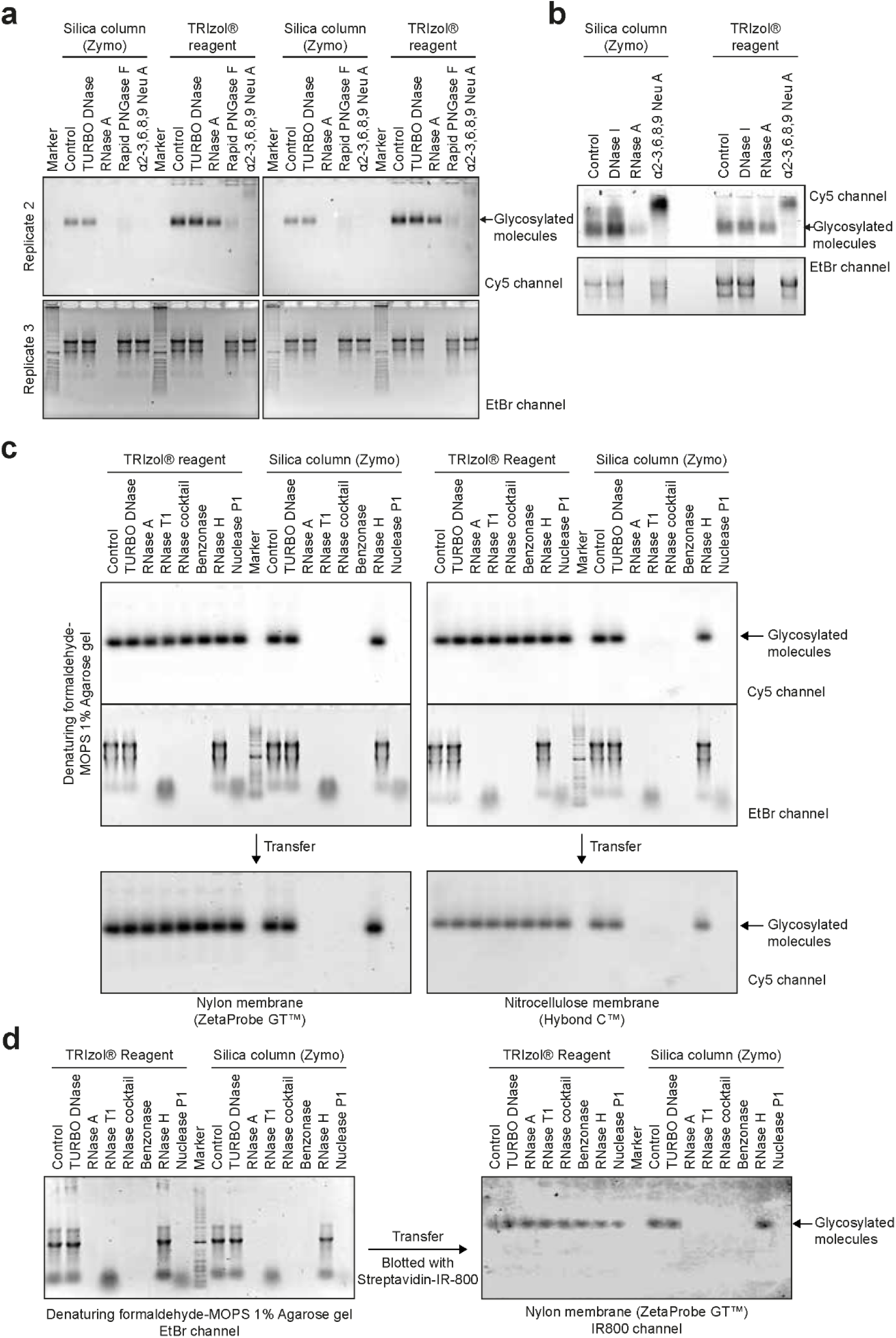
Comparison between silica-based column and TRIzol purification for the last clean-up step, related to Figure 2. **a.** Data represent other two replicates done in Fig. 2d. **b.** Independent reproduction from a different lab’s experiments for the effect of TRI Reagent and silica column purification on recovery. Final RNA purification in the silica column was performed at 60% EtOH. DNase I was used as an alternative for DNA degradation. **c.** Glycan detection after various nucleases. **d.** Blotting of biotinylated glycans using streptavidin-conjugated IR800 dyes on the nylon membrane. Right, gel image of EtBr-stained RNA samples. Left, fluorescent image using IR800-streptavidin in 800-nm channel.

**Extended Data Fig. 3.**
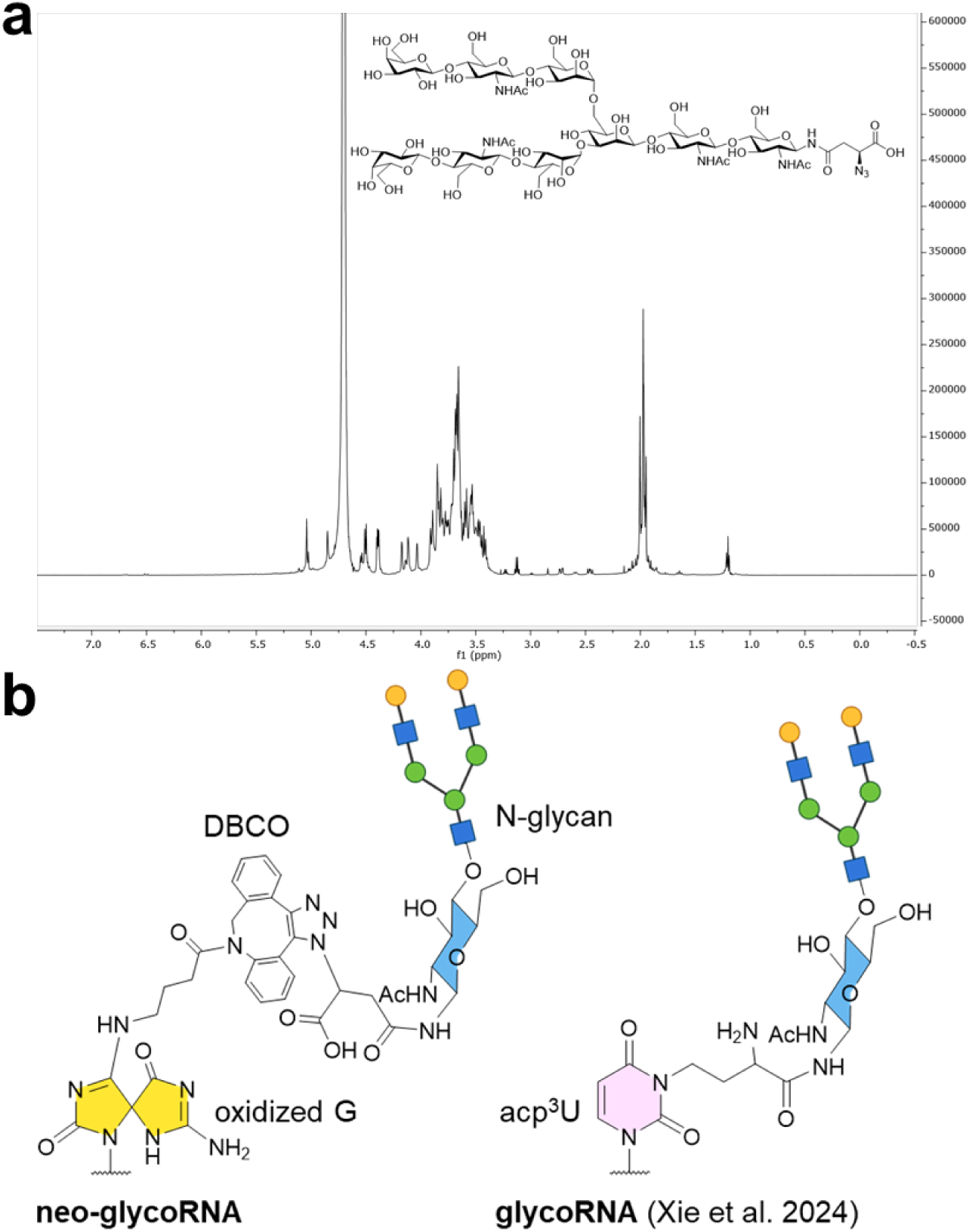
Structure and supporting data for neo-glycoRNA. **a.** Proton nuclear magnetic resonance demonstrating purity and quality of the azide-functionalized N-glycan. **b.** Chemical structure of neo-glycoRNA and the comparison with reported glycoRNA. Abbreviations: G, guanosine; acp^3^U, 3-(3-amino-3-carboxypropyl)uridine.

**Extended Data Fig. 4.**
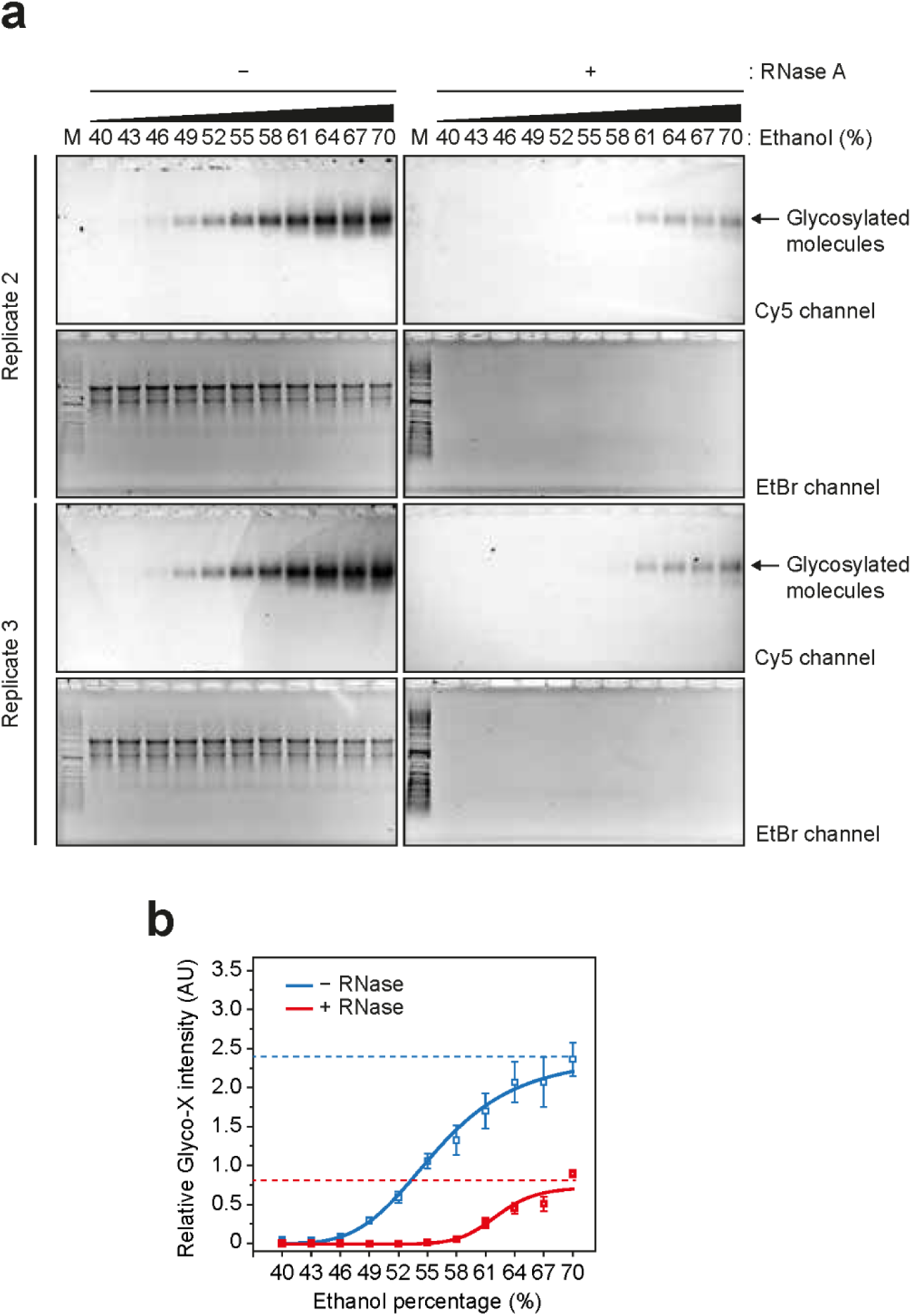
Recovery of glycosylated molecules at various ethanol percentages, related to. **Figure 3** **a.** Data represent other two replicates done in Fig. 3c. **b.** Relative glycan intensity calculated from the data points in Fig. 3c and Extended Data Fig. 3a. Error bars represent s.d.

**Extended Data Fig. 5.**
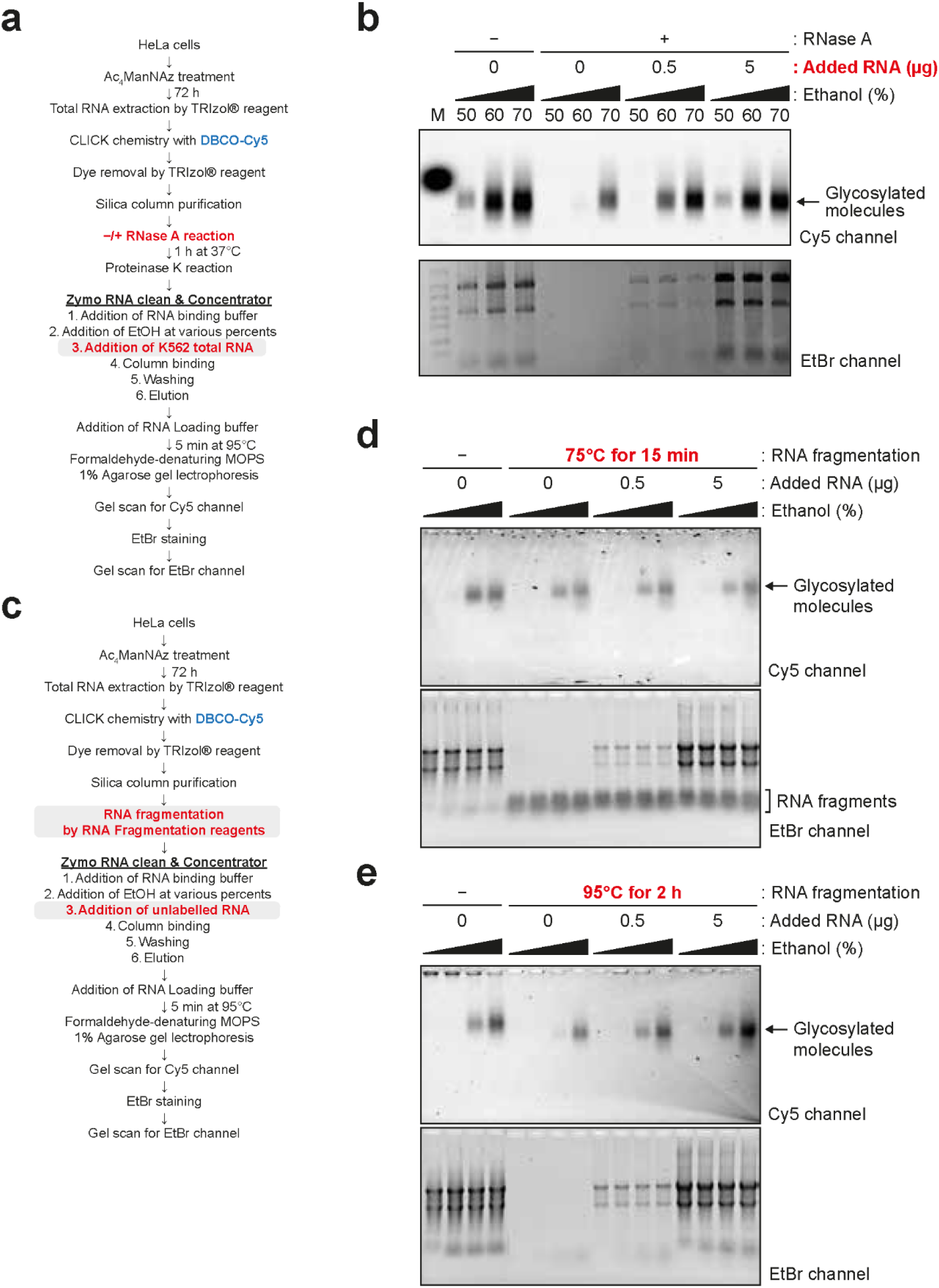
Added RNA but not DNA improves the recovery rate of glycosylated molecules in RNA-depleted conditions, related to Figure 5. **a.** Schematic for the experiment in Extended Data Fig. 5b. **b.** Data from the Li’s experiment. Total RNA extracted from K562 cells was used as an alternative for added HeLa total RNA. **c.** Schematic for experiments in Extended Data Fig. 5d and 5e. **d.** Glycan recovery by the added RNA in the mild RNA fragmentation condition. Partially fragmented RNAs are indicated in EtBr channel. **e.** Glycan recovery by the added RNA in the complete RNA fragmentation condition.

